# The circulatory physiopathology of human red blood cells investigated with a multiplatform model of cellular homeostasis. III. Senescence changes during the full circulatory lifespan

**DOI:** 10.1101/2020.03.07.981803

**Authors:** Simon Rogers, Virgilio L. Lew

## Abstract

Human red blood cells (RBCs) have a circulatory lifespan of about four months. Under constant oxidative and mechanical stress, but devoid of organelles and deprived of biosynthetic capacity for protein renewal, RBCs undergo substantial homeostatic changes, progressive densification followed by late density reversal among others, changes assumed to have been harnessed by evolution to sustain the rheological competence of the RBCs for as long as possible. The unknown mechanisms by which this is achieved are the subject of this investigation. Each RBC traverses capillaries between 1000 and 2000 times per day, roughly one transit per minute, a total of about 2•10^5^ transits during their lifespan. A dedicated Lifespan model of RBC homeostasis was developed as an extension of the RCM introduced in the first paper of this series to explore the cumulative patterns predicted for repetitive capillary transits over a standardized lifespan period of 120 days, using experimental data to constrain the parameter space. Capillary transits were simulated by periods of elevated cell/medium volume ratios and by transient deformation-induced permeability changes attributed to PIEZO1 channel mediation as outlined in the second paper of this series. The first unexpected finding was that quantal changes generated during single capillary transits cease accumulating after a few days and cannot account for the observed progressive densification of RBCs on their own, thus ruling out the quantal hypothesis. The second unexpected finding was that the documented patterns of RBC densification and late reversal could only be emulated by the implementation of a strict time-course of decay in the activities of the calcium and Na/K pumps, but only in addition to the quantal changes. These results showed that both quantal changes and pump-decay regimes were necessary to account for the documented lifespan pattern, neither sufficient on their own. They also suggested a strong selective component in the pump decay sequence. A third finding was that RBCs exposed to levels of calcium permeation above certain thresholds in the circulation could develop a pattern of late or early hyperdense collapse followed by delayed density reversal. When tested over much reduced lifespan periods the results emulated the known circulatory fate of irreversible sickle cells, the cell subpopulation responsible for vaso-occlusion and for most of the clinical manifestations of sickle cell disease. Analysis of the results provided an insightful new understanding of the mechanisms driving the changes in RBC homeostasis during circulatory aging in health and disease.

## Introduction

Bridging the information gained from the study of single capillary transits [previous paper] to account for the changes documented for full lifespan periods [1] proved a formidable challenge. Initial attempts at approaching this investigation using the red cell model [2, 3] with a protocol of multiple repetitive extensions of dynamic state pages, with the level of accuracy demanded by the infinitesimal magnitude of each single transit [3], led to model stagnation after only a day or two, thus disabling this brute force approach. It became clear that to undertake a proper lifespan study it was necessary to reset the red cell model into a new framework specifically dedicated to follow the homeostatic changes of RBCs throughout their long circulatory journey.

The lifespan model introduced here was developed by one of us (SR) to enable a detailed exploration of the multiple factors shaping the changes RBCs experience in the circulation and to explain the mechanisms at work. The results showed that transient increases in PIEZO1-attributted calcium and anion permeabilities during capillary transits were necessary but not sufficient for generating gradual densification. For gradual densification and late density reversal timed decays in the activities of the calcium and Na/K pumps proved absolutely necessary. These results allowed the formulation of a new mechanism of RBC senescence combining both quantal and pump-decay processes, compatible with all available evidence. The simulations confirmed the versatility and potential of the Lifespan model as a dedicated tool for the investigation of the mechanisms controlling the hydration state of RBCs during circulatory aging in health and disease.

## Methods

### Open access to the Lifespan model

As for the core RCM model [2, 3], the Lifespan model, Lifespan*.jar, is available for downloading from the GitHub repository https://github.com/sdrogers/redcellmodeljava. The model operates as a *.jar programme within the JAVA environment which needs to be preinstalled. It is recommended not to alter the original file name as it contains coded information on date and update status. Altered names are best applied to shortcuts. Double-click to activate the programme. Operation of the RCM model generates *.csv files containing the results of simulations with identical format to the one generated by the RCM programme. By default, these are saved within the same directory as the one from which the programme was operated, a choice easily modified from within the programme. The modified defaults set on the model offered for download, if left unchanged, will automatically run the 120 lifespan trajectory of a RBC used as the “reference” pattern in the current study, as reported by the black curves in figures 3-5 and 7-9.

**Figure 1.**
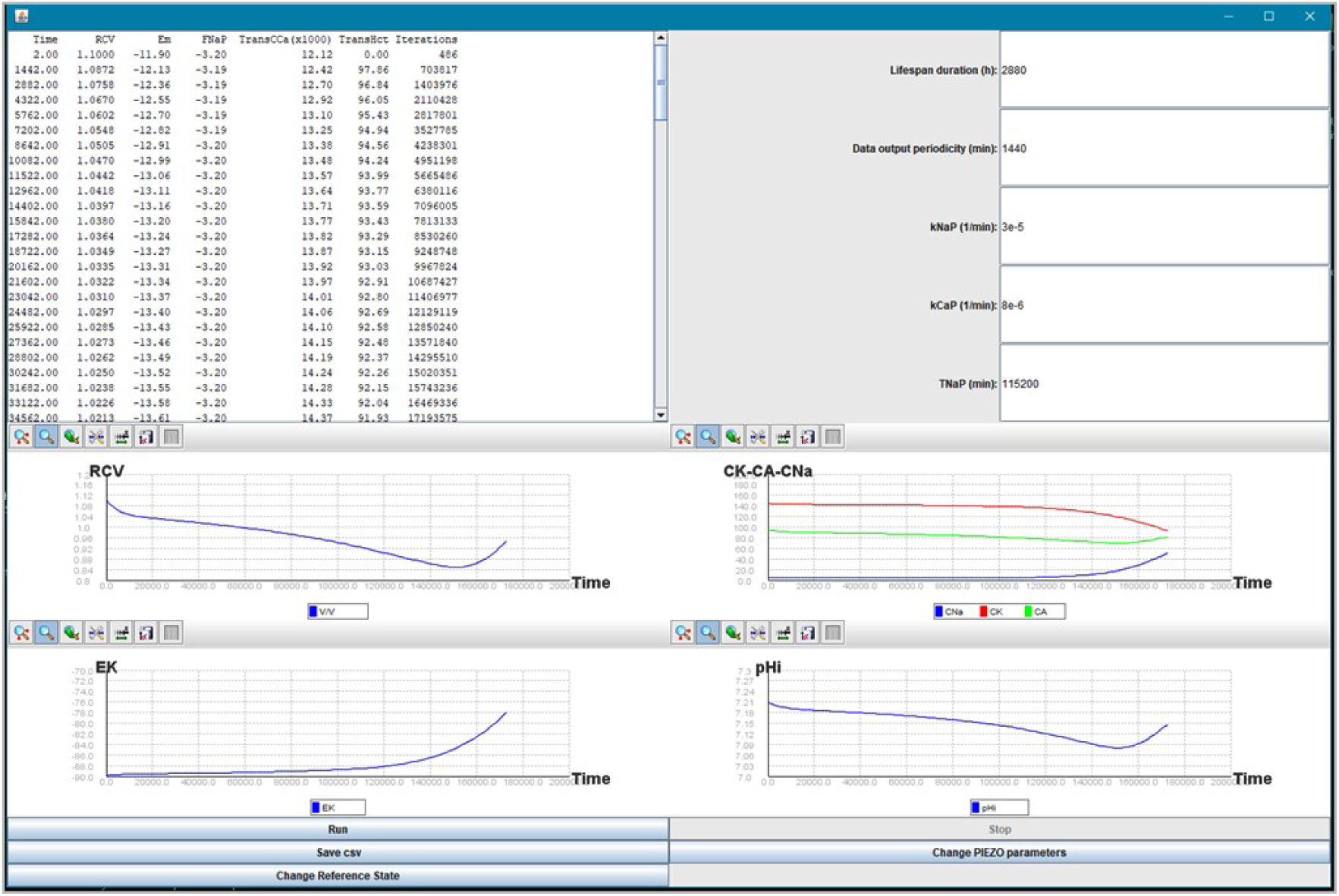
User interface of the Lifespan model. The interface was designed to offer the user information on the go during the computations allowing the model to be stopped, the protocol to be modified and the programme to be run again from the start. The “Change Reference State” tag calls up a window with options for controlling the initial condition of the cell; the “Change PIEZO parameters” tag brings up the window for setting the PIEZO1-mediated permeabilities and the duration of the open-state. Three additional tags implement instructions for running (Run) and stopping (Stop) the computations and for generating csv files at the end of a run (Save csv) with the results displayed in the same format as that described before [2]. The main user interface is divided in three main panels: the top-right panel lists five time-dependent values with the defaulted unit dimensions found most convenient during exploratory preliminary work; from top to bottom: Lifespan duration (h), data-output periodicity (min), rate constants of exponential decay for the Na/K (kNaP) and calcium pumps (kCaP; 1/min), and delayed onset time for Na/K pump decay (TNaP; min). In the running of the Lifespan model, selected data appear listed in the top left panel: Time (in min), Relative cell volume (RCV), Em (mV), Na/K pump-mediated Na efflux (FNaP, mmol/Loch), Trans-CCa (μmol/Loc), Trans-Hct (%), and number of model cycles (Iterations) in between data points. Trans-labelled variables report values at the end of the last capillary transit for the time point listed under Time. Selected variables are shown in graphic format in the bottom panel, clockwise from top left: RCV, CNa & CK & CA (mmol/Loc), pHi, and EK (the potassium equilibrium potential, mV). All default values for parameters and initial variables correspond to those used for the pattern defined by the Reference lifespan curve (text and Fig 2). It was necessary to generate two csv results files because of differences in the values of certain variables if recorded at the end of transit (Transit-named csv file) or at the end of intertransit periods for each time point. The differences concerned mainly the following variables: CVF/Ht, CCa, CCa2+, CH, MNa, MK, MA, MH, FCaP, FKGardos and all FzX.

**Figure 2.**
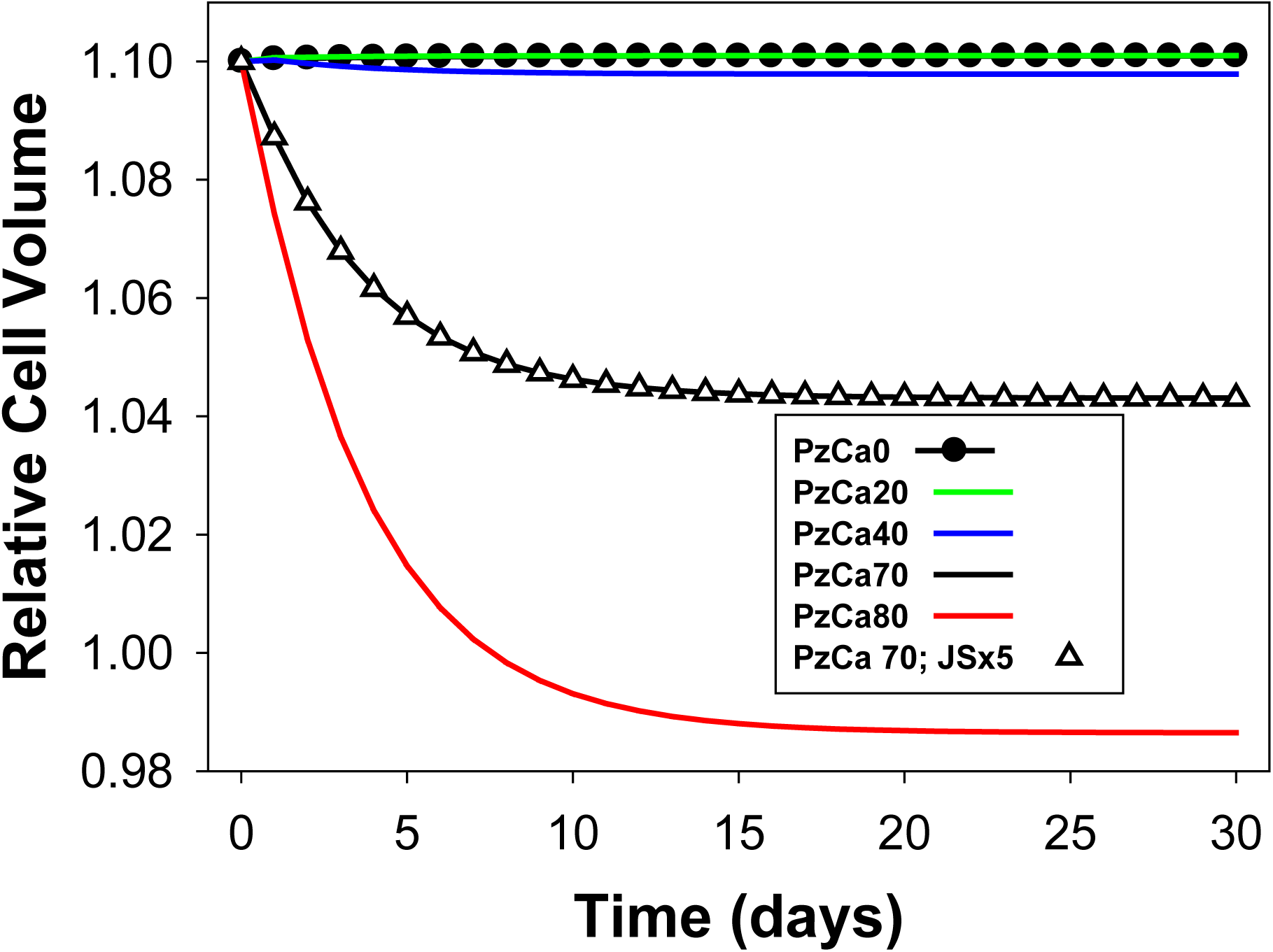
Predicted lifespan changes in Relative Cell Volumes at different PzCa levels. The trend shown only for the first thirty days remained unchanged for 120 days. Pump decay rates were set at zero. Triangles: effects of increasing JS turnover rate five-fold at PzCa = 70/h. Note that even at the highest PzCa shown, final dehydration does not exceed 10% of initial RCV. Except for initial periods of 10-20 days further progressive dehydration throughout RBC lifespans cannot be induced by the cumulative effect of episodic PIEZO1 activations during capillary transits.

**Figure 3.**
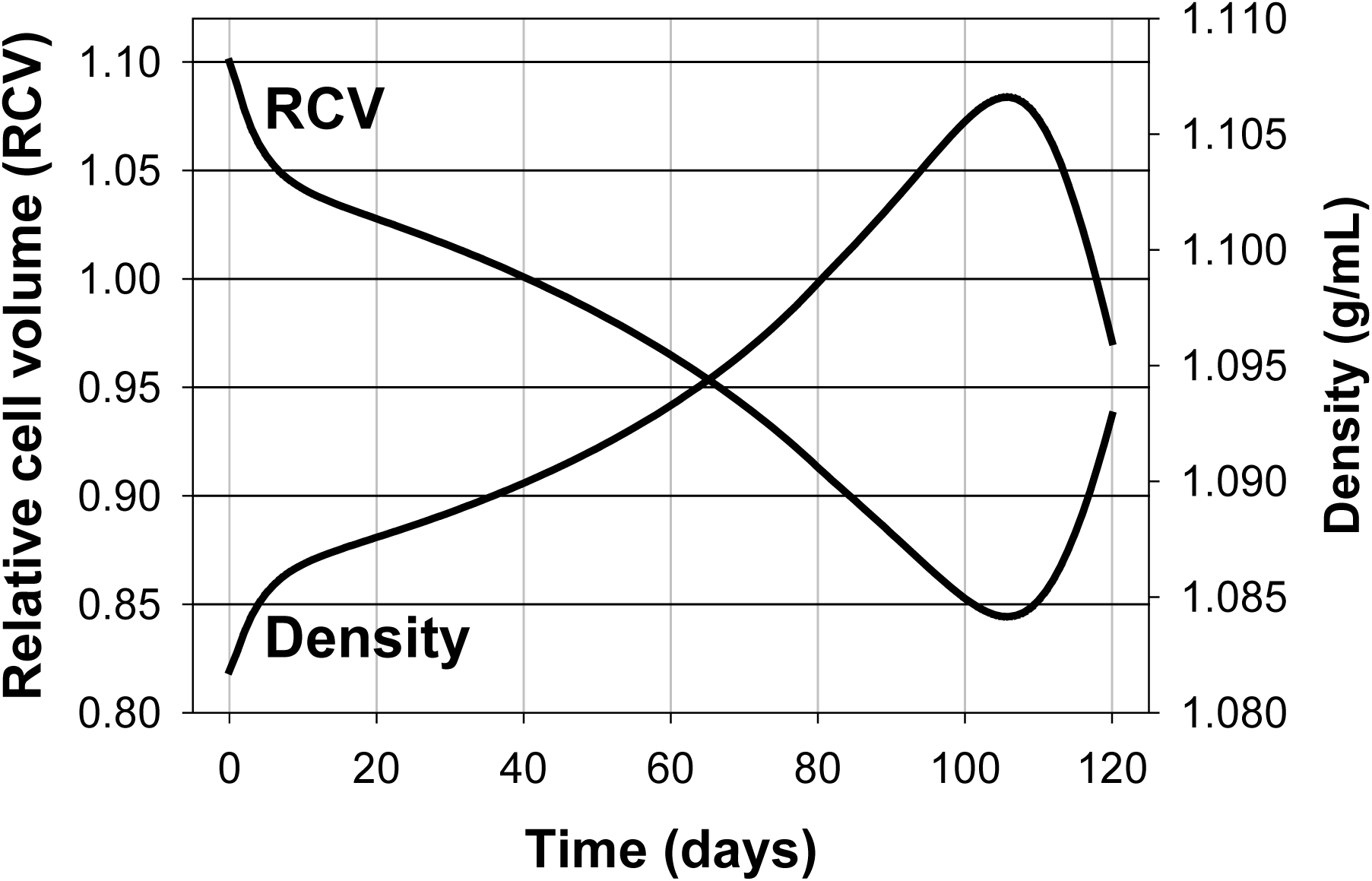
Predicted patterns of change in relative cell volume and density over a standardized 120 day lifespan period. The lines join data points collected at daily intervals. Simulations run with the cell volume fraction set at 0.9 during the brief capillary transit period [3]. The “Restore medium” was set to YES to prevent carryover changes in medium concentrations during inter-transit periods. **RCV:** Relative cell volume, a convention adopted in RBC homeostasis models to report RBV volumes relative to a standardized value of 1 L/Loc attributed to a RBC defined with 0.75 Lcw/Loc, 0.25 LHb/Loc and 340 gHb/Loc ([2], Appendix). **Density:** Density profile, in g/ml. The initial condition of the cell was defined with Vw of 0.85 Lcw/Loc, CNa of 5 mM, CK of 145 mM, and a Na/K pump-mediated Na^+^ efflux rate of −3.2 mmol/Loch, an approximate representation of the condition of a recently transitioned cell from reticulocyte to mature RBC with its full complement of haemoglobin set at 340 gHb/Loc. The parameter values used were: OS, 0.4s; PzCa, 70/h; PzA, 50/h (no significant differences between 30/h and 50/h for PzA, the range observed under on-cell patch clamp [19]); kCaP, 8e-6/min; kNaP, 3e-5/min; TNaP, 115200min; PzNa and PzK were set to zero. The RCV curve is used as a standardized reference (Ref) for analysing the effects of parameter variations in figures 4 and 5.

**Figure 4.**
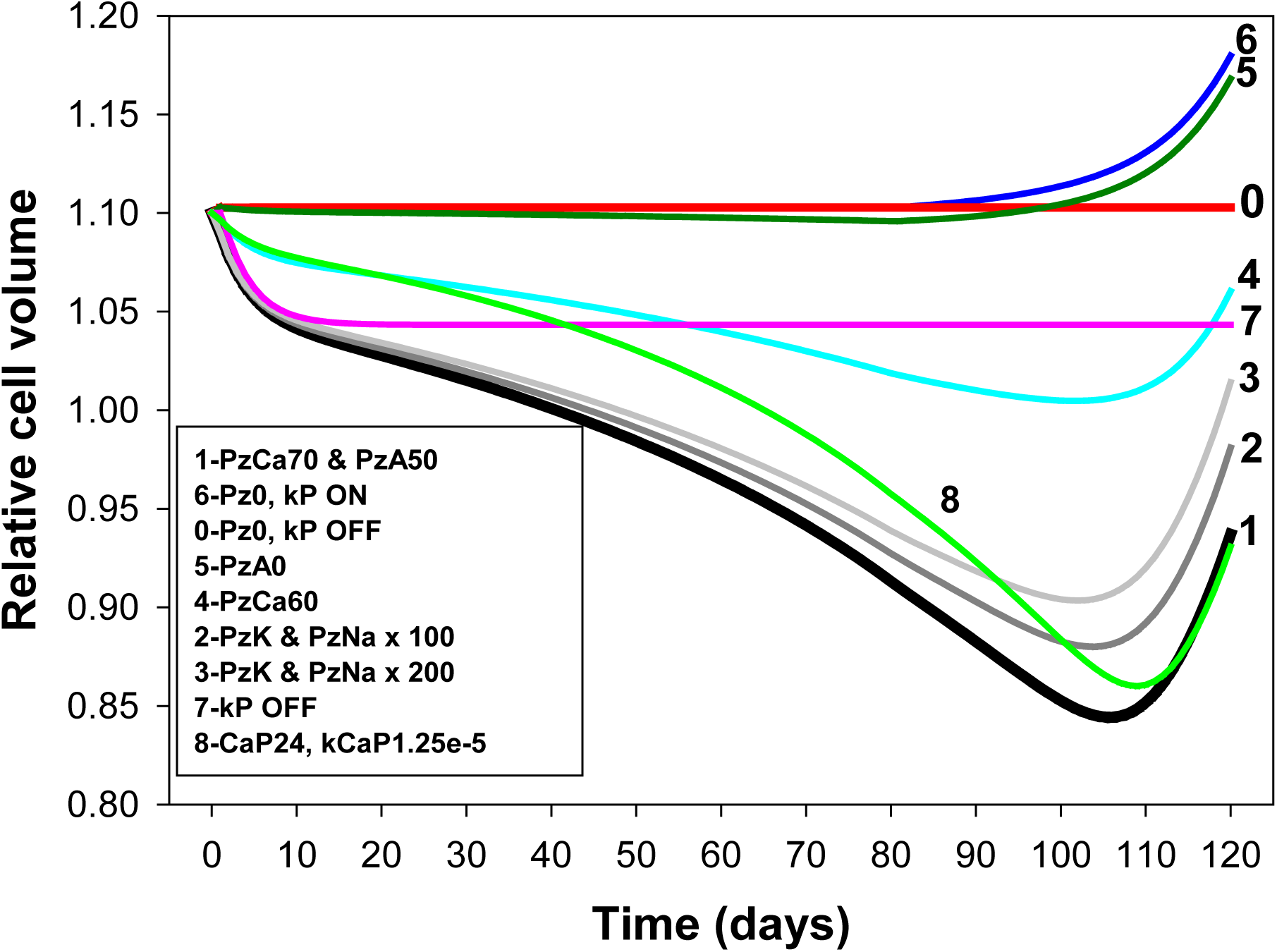
Effect of changes in PIEZO1-mediated ionic permeabilities and pump decay rates on the lifetime pattern of cell volume change. **1**. The **p**arameter values for Reference curve 1 (black) are as reported in the legend of Fig 2. **0:** With PzX = 0 and no pump decay the model computes a flat response over the full 120 days period demonstrating the robust stability of the Lifetime computations. **7:** With the PzX set as for the reference curve (curve 1) but with no pump decay (curve 7, kP OFF) there is no progressive dehydration-densification, only the early quantal dehydration reported in Fig 2. **6:** With PzX = 0 and pump decays set to ON (curve 6, Pz0, kP ON) there is no dehydration phase, only late hydration following delayed Na/K pump decay. **5:** Same as curve 1 but with PzA set to zero showing how extremely limiting the anion permeability can be to both initial and cumulative dehydration responses. **4:** Relatively minor reductions in PzCa from 70/h (Curve 1) to 60/h (curve 4) reduce initial and cumulative dehydration responses outside observed ranges. **2 & 3:** Large changes in PIEZO1-mediated Na^+^ and K^+^ permeabilities, curves 2 and 3, have relatively minor effects, mostly on the timing and magnitude of the late density reversal response, rendering the Lifespan model a poor predictor of their likely real values. **8:** Protocol identical to that of reference curve 1 but for a cell defined in the RS with a FCaPmax of 24 instead of 12 mmol/Loch; the increased pump strength reduced the extent of early dehydration and in order to approximate the reference dehydration pattern as shown it was necessary to increase kCaP from 8e-6 to 1.25e-5 min^-1^.

**Figure 5.**
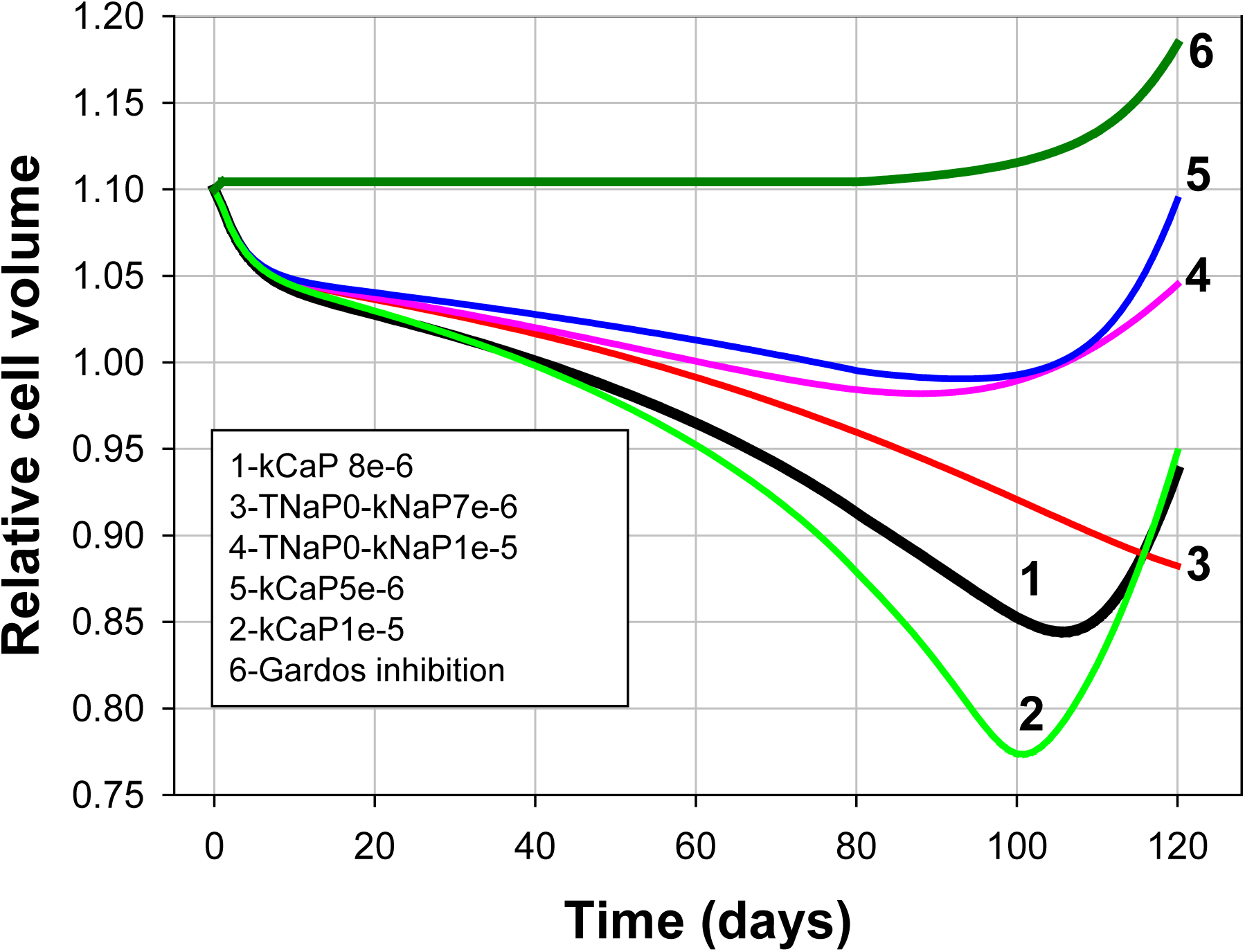
Effects of pump decay rates and timings, and of Gardos channel inhibition on the predicted lifetime pattern of cell volume change. Parameter changes are listed relative to reference curve 1. For the simulation shown in curve 6 the Gardos channel Fmax was changed from 30/h to zero in the Reference State. For curves 3 and 4 TNaP was set to zero and the decay rates to 7e-6/min and 1e-5, respectively. For curves 2 and 5 PMCA decay rates were set to 1e-5 and 5e-6, respectively.

**Figure 6.**
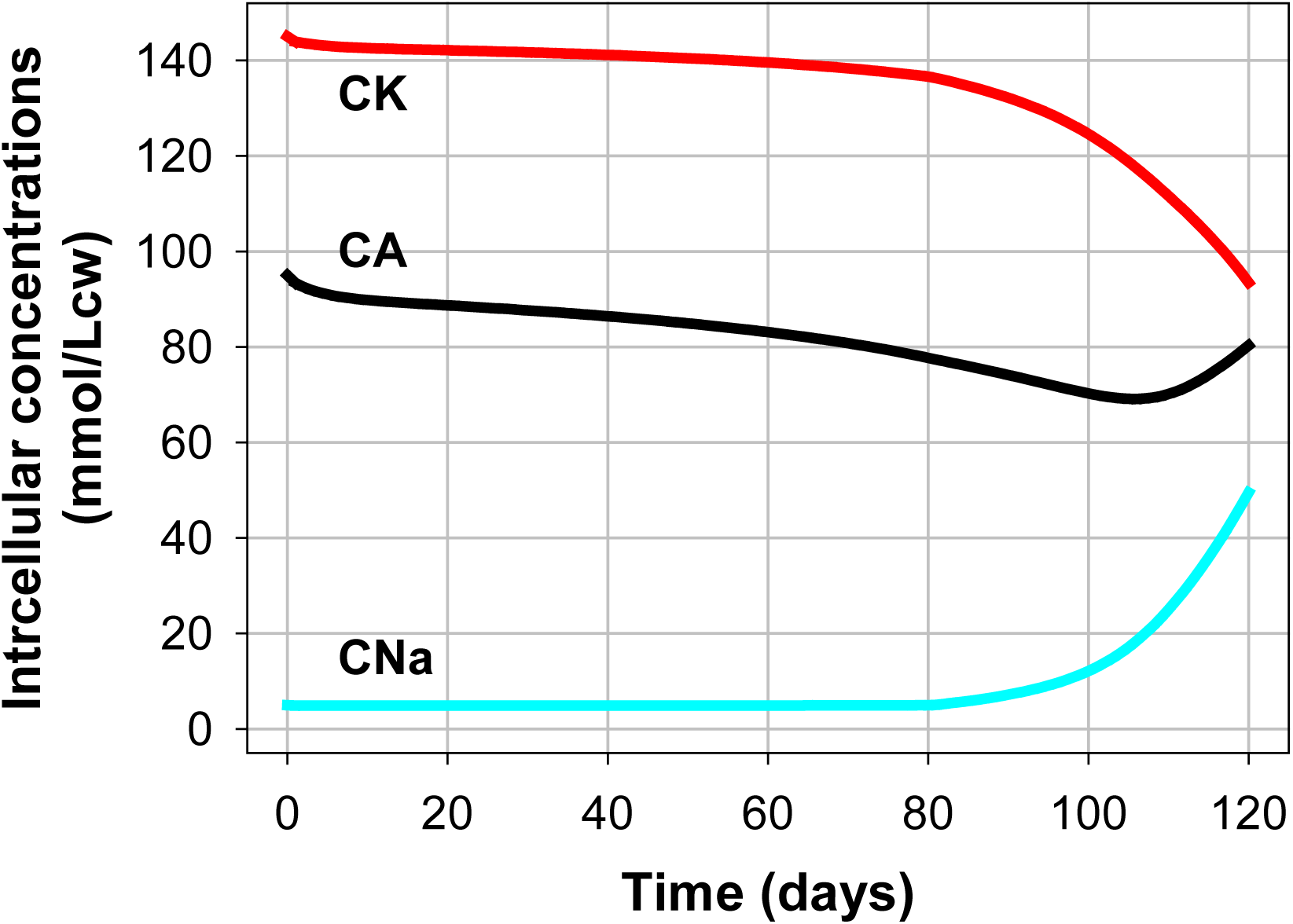
Predicted lifespan changes in the intracellular concentrations of Na (CNa), of K (CK) and of diffusible anions A (CA) for the conditions of the reference pattern (Fig 2). Note that because PzNa and PzK were set at zero in the minimalistic representation of the reference pattern, the predicted changes shown in this figure result solely from the cumulative effects of periodic Gardos channel activation and Na/K pump decay.

**Figure 7.**
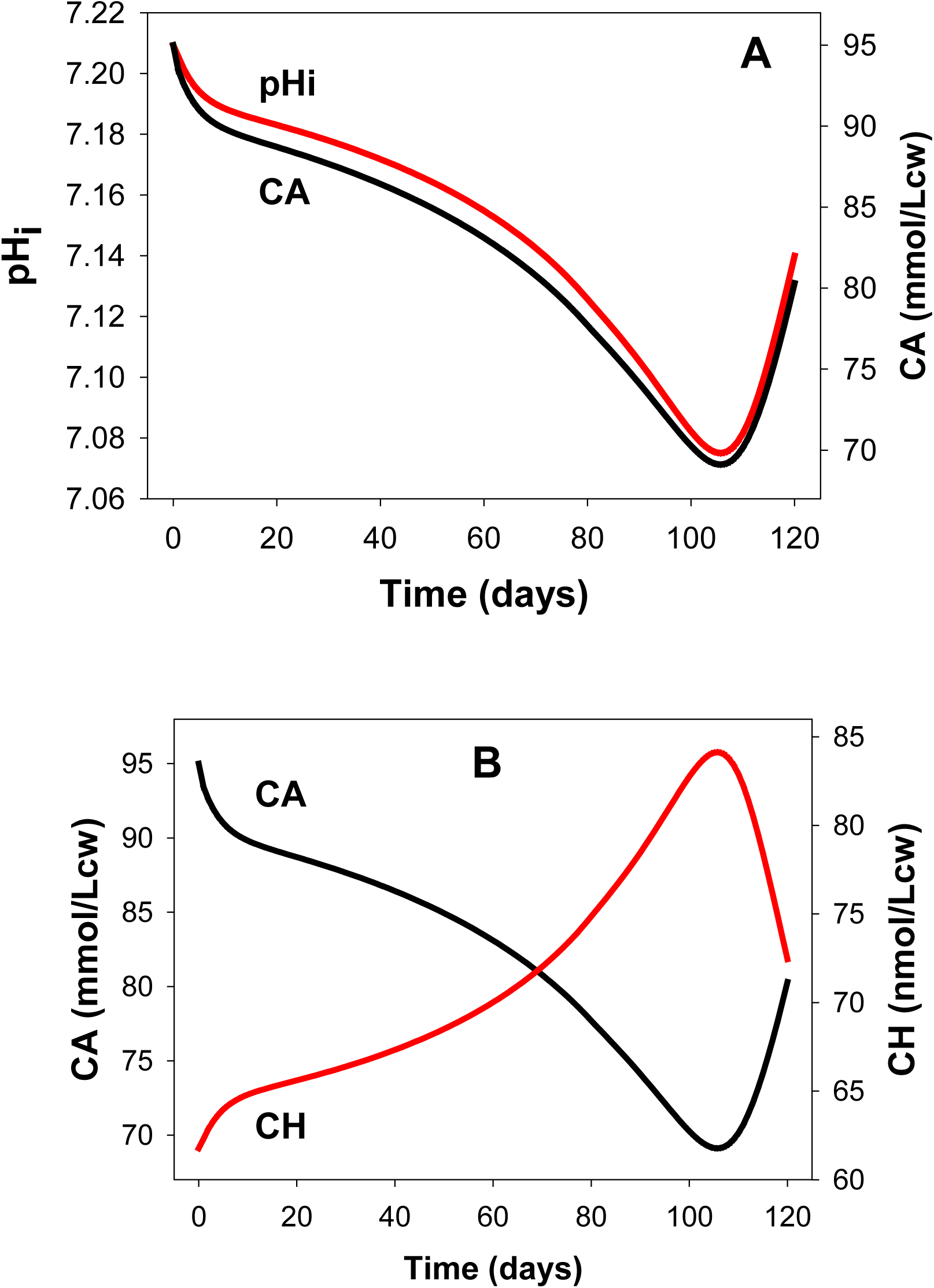
Predicted lifespan changes in cell pH (pHi) and in the intracellular concentrations of A^-^ (CA) and of H^+^ (CH) for the conditions of the Reference pattern (Fig 2). Besides minor contributions from variations in haemoglobin buffering, most changes in CH are driven primarily by changes in the anion concentration gradient (rA) acting through the Jacob-Stewart mechanism causing rH to approach rA during periods between capillary transits []. At constant MA all anion gradient changes apply to CA, thus generating similar time-courses for CA and pHi (**Panel A**), or mirror image changes for CA and CH (**Panel B**).

**Figure 8.**
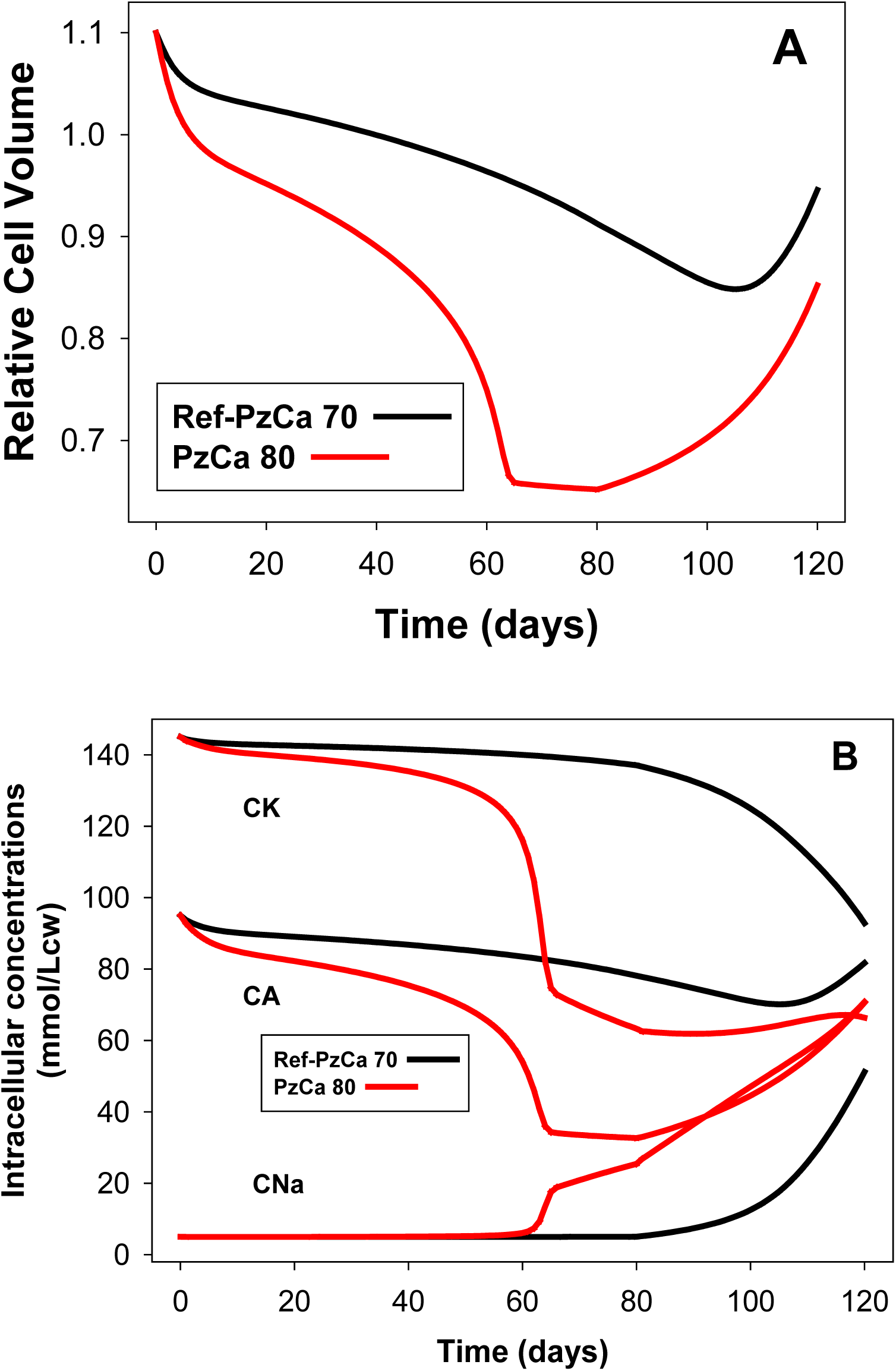
Predicted changes in RBC volume (A), and in cell concentrations (B) of K(CK), of Na(CNa), and of diffusible anion(CA) during hyperdense collapse induced by elevated PzCa. The changes induced by setting PzCa = 80/h (red curves) are compared with those of the reference pattern with PzCa = 70/h (black curves).

**Figure 9.**
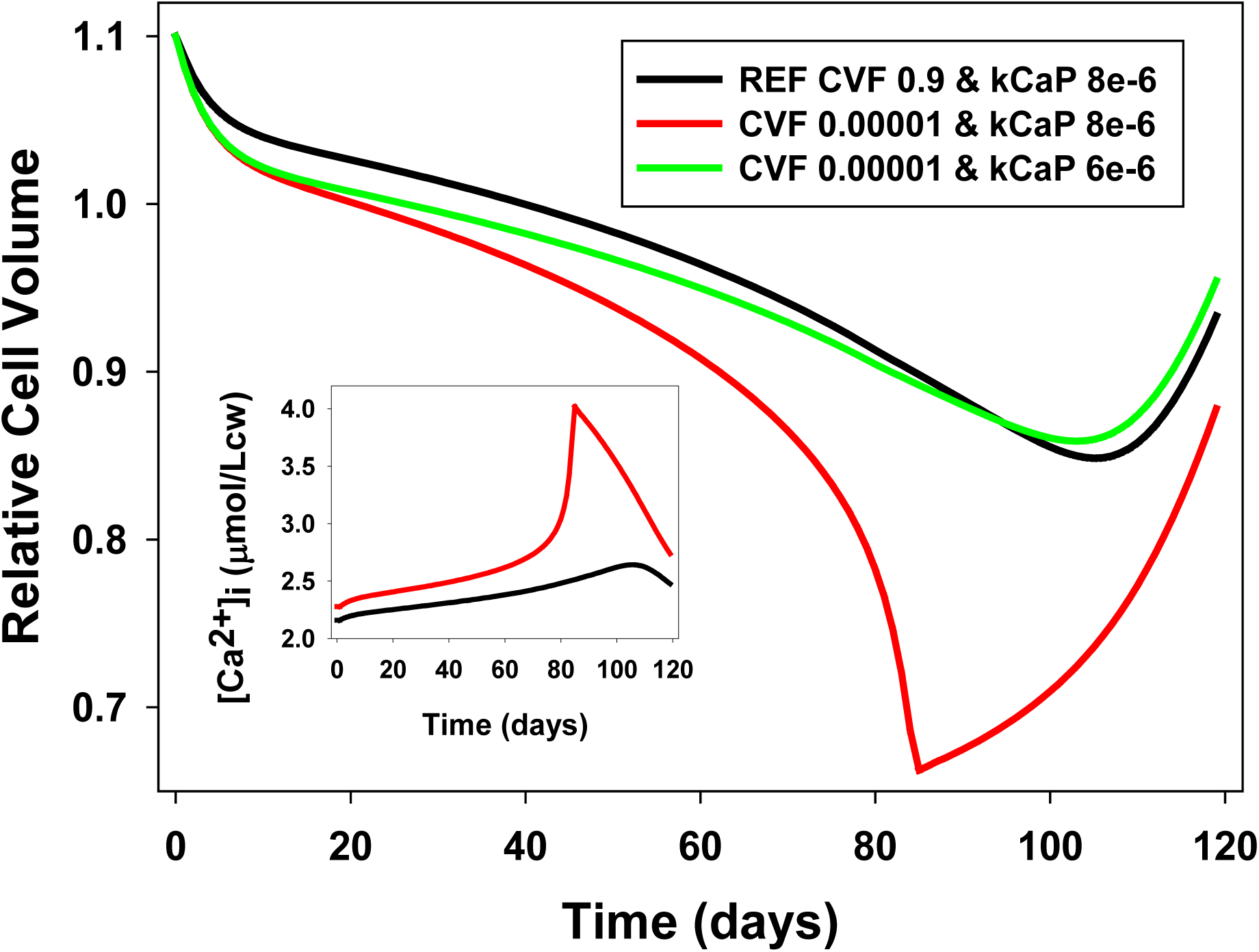
Effects of high and low cell volume fractions (CVF) during capillary transits on the lifespan patterns of changes in cell volume and [Ca^2+^]_i_. Reference pattern (black) modelled with CVF of 0.9 during capillary transits. Test curves (red and green) modelled with CVF of 0.00001 throughout transit-intertransit periods. PMCA decay rates (kCaP), as indicated in the figure (in min^-1^). Note how the reduction in kCaP from 8e-6 (red) to 6e-6 (green) prevents hyperdense collapse at CVF 0.00001 and restores pattern towards Reference curve (black). **Inset:** [Ca^2+^]_i_ values recorded at the end of PIEZO1 open states on the last model iteration of each day (col 19 of the transit csv files). Note the increasing divergence between REF (black) and low-CVF (red) curves leading to hyperdense collapse. [Ca^2+^]_i_ late reversal results from terminal rehydration of the cells.

### Modelling the changes in RBC homeostasis during a standardized 120 day lifespan

The red cell model, framed into the new Lifespan model (Fig 1; details in the legend of Fig 1) allows testing a vast range of protocols using a dedicated user friendly interface. The operator sets the initial condition of the RBC, the parameter values under test, the duration of the simulation, the frequency of data outputs, and runs the model. While running, the user interface displays the changes in time of selected system variables in numerical and graphic formats for the full duration of the simulation. At the end of each simulation the full data sets can be saved in csv-file format. Consistent with the strategy followed in all previous RBC model versions, the changing “Relative cell volume” of modelled RBCs is computed by reference to a cell defined with 0.75 Lcw/Loc (litres of cell water per litre original cells), 0.25 LHb/Loc (litres of haemoglobin per litre original cells, equivalent to 340g Hb with a specific density of 1.36g/mL [4]), and a density of 1.090g/mL, values very near measured means in samples from healthy adult subjects.

The most relevant and widely supported experimental constraint to guide the model outcomes concerns the age-dependent progressive densification of RBCs. Recently matured RBCs with their full complement of haemoglobin are found at densities about 1.080-1.085 g/ml, whereas aging RBCs are seldom found beyond densities of 1.105-1.110 g/ml [5-9] These boundaries then define a working range of age-related density progression in human RBCs from healthy donors for most of the RBC lifespan. A second important constraint for the model to comply with is the late density reversal phenomenon, originally reported by Borun et al., [5], overlooked for decades, and finally confirmed and extended in recent research [9-11]. Borun et al., [5] showed that between about 70 and 100 days in the circulation RBCs reverse their densification pattern, undergoing a partial, late and terminal density reversal. Progressive densification within established boundaries and late density reversal will be applied as the prime constraints on the parameter set for the model outcomes to comply with. Additional compliances with available evidence will be considered along the different stages of this investigation.

## Results and Analysis

The first simulations with the lifespan model were carried out to investigate whether the documented patterns of progressive dehydration-densification could be fulfilled by the cumulative effects of single transits, as suggested by the quantal hypothesis. Figure 2 shows the predicted results at different PzCa levels. The levelled off trend shown for the first thirty days continued unchanged for 120 days (not shown). It can be seen that RBC densification increases with PzCa but only for the first 10-20 days, levelling off thereafter without any further progression. Even at PzCa = 80, RBC density did not progress beyond the 1.090g/mL level. Deep data analysis of the variables in the csv files at high PzCa shows that during the gradual dehydration of the first days single passages contribute diminishing quantal dehydration steps slowly progressing towards a homeostatic condition of fluid balance. The initial fluid gained on PIEZO1 activation during each transit gradually becomes balanced by dehydration towards the end of the intertransit stage precluding any further densification. In terms of the transport mechanisms involved, using the representations in Fig 4 of the previous paper [3], this approach to balance implies that |W1| becomes progressively equal to |W2+W3+W4|, i.e. fluid influx through PIEZO1 (W1) becoming gradually closer to the sum of fluid losses through Gardos channels (W2), PMCA (W3) and JS (W4) in each transit-intertransit period. This result therefore rules out lifespan mechanisms based *exclusively* on cumulative effects of individual quantal changes during capillary passages. To generate a pattern in full compliance with the densification-reversal sequence, the search focused on transport systems with well documented patterns of activity decline with RBC age, the calcium and sodium pumps [10, 12-18].

Measured PMCA Fmax distributions in RBCs from five difference donors [12] rendered coefficients of variation (CV) of between 45 and 53%, with means to the right of the median in each, right skews ranging from 1.40 to 1.67. To represent such a decay process in the model we considered linear and exponential decay functions. The symmetry of linear decay distributions was incompatible with the observed skewed pattern. Exponential decay functions on the other hand generate right skews and were the obvious choice. There is no such detailed information on the modality of sodium pump decay. The few reliable findings document differences between light and dense cell fractions, substantial declines in [H3]-ouabain binding and ouabain-sensitive potassium transport in the dense cell fractions [15, 18]. On this background, a large number of preliminary tests were carried out with the Lifespan model seeking to establish whether it was possible to generate a pattern in full compliance with the known densification-reversal sequence. Unexpectedly, the pattern that emerged from these trials and that fulfilled the expected compliance could be implemented with a single and very restricted set of parameter values.

### The Reference pattern

In order to report the process followed to arrive at this result in the simplest possible terms we generated a protocol with default values which provides a standardized reference curve, the Reference pattern (Fig 3), that can be used for comparison with the effects of parameter variations thus allowing a detailed analysis of the mechanisms involved (Figs 4 and 5).

Figure 3 shows the predicted changes in relative RBC volume and density in simulations aimed at defining the minimal set of parameter values able to provide full compliance with the experimental constraints. These values are listed in the legend of Fig 3. As explained in detail for single transits, realistic representations required high values for the cell/medium volume ratio (equivalent to high haematocrits) during capillary transits. All the simulations reported here were set with an open state (OS) duration of 0.4s and with one minute duration for inter-transit periods. For an OS of 0.4s a nominal calcium permeability (PzCa) through PIEZO1 channels of 70h^-1^ rendered the results shown in Fig 3 for our reference condition.

The volume curve (Fig 3) will be used as our reference for all further explorations of the parameter space (black curves in Figs 3-5 and 8). The volume and density curves (Fig 3) are mirror images of each other as expected from the model-implemented assumption that the haemoglobin complement of each cell after maturation from the reticulocyte stage remains constant throughout its lifespan. Thus, changes in density reflect only variations in hydration state. The rate of volume change during the densification phase, between days 10 and 100 was about −0.2%/day (Fig 3).

The mechanism by which gradual decay in the activity of the calcium pump induces progressive dehydration operates by gradually weakening the calcium pump and so delaying the extrusion of the Ca^2+^ gained during each PIEZO1 open state (Fig 4C). This delay extends the duration of elevated [Ca^2+^]_i_ states in successive capillary transits progressively increasing the extent of quantal dehydration ([3], Figs 5C-E). This breaks the balance which halted dehydration progress during consecutive quantal transits under the assumption of constant pump strength in the quantal hypothesis (Fig 2). PMCA decay changes the quantal hypothesis stalemate from |W1| = |W2+W3+W4| to an increasing imbalance where |W2+W3+W4| > |W1| ([3], Fig 4).

The exponential decay rate of the PMCA found to comply best with observed densification patterns (Fig 3) predicted an age-determined distribution of PMCA Fmax activities for circulating RBCs in remarkable agreement with experimental results [12]. Measured PMCA Fmax distributions in RBCs from six difference donors rendered coefficients of variation (CV) of between 45 and 53%, with means to the right of the median in each. The PMCA decay function that offered the best fit to observed densification patterns in our model simulations (Fig 3) followed the equation y = y°*exp(-(8*10^−6^)t). The y-values computed from this equation for the 120 days of our standard lifespan period, report the age–dependent Fmax decline of the model PMCA with a distribution whose statistical parameters can be compared to those of measured ones. Using the default PMCA Fmax value of 12 mmol/Loch for y° the resulting statistical parameters for the Ref PMCA decline function were: median, 6.12; mean, 6.64; SD, 2.69; CV 46% and skew of 1.2 which compares with measured skews of between 1.4 and 1.7 in RBCs from six different donors. Similar CV and skew values are obtained with different y° and kCaP pairs constrained to approximate the reference densification pattern, as for curve 8 in Fig 4, for instance. The similarity between measured and predicted CVs and skews suggests that the variation in PMCA activity in RBC populations from healthy adult donors is almost entirely age-related, and that its decay pattern was harnessed by evolution to extend the circulatory lifespan of RBCs.

As analysed in detail below, a delayed onset of an exponential decay in sodium pump activity was required for early permissive densification and late density reversal.

### Analysis of the effects of PIEZO1-mediated permeabilities and pump decay

When all PIEZO1-mediated permeabilities and pump decay rates are set to zero (Fig 4, curve 0) the model computes a flat response over the full 120 day period and 10^9^ model iterations demonstrating the noise-free robust stability of the Lifetime computations. Restoring pump-decay rates with PIEZO1 blocked (Fig 4, curve 6) retains the flat response until a late hydration-densification stage resulting from delayed Na/K pump decay. This response also demonstrates the absolute need to involve PIEZO1 mediation in the generation of the reference pattern, pump decay on its own does not generate gradual densification indicating that sufficient Ca^2+^ extrusion capacity remains to effectively balance Ca^2+^ influx through the intrinsic Ca^2+^ permeability of the RBC membrane for the whole lifespan of the cells.. With pump decay rates set to zero (Fig 4, curve 7), the model predicts a minor dehydration over the first few days, followed by a flat volume response, repeating conditions analysed for Fig 2 in the context of all other tests. Gradual decays in the activities of the calcium and Na/K pumps are therefore a necessary condition for generating the pattern of progressive changes in RBC volume during circulatory senescence (Fig 3). Taken together (Figs, 2-4), these results show that both processes, quantal dehydrations and pump decays are necessary for shaping the observed pattern of lifespan RBC volume changes, neither sufficient on its own.

We investigate next the role of the anion permeability in the generation of the lifetime dehydration-densification pattern. A substantial increase in anion permeability had been detected associated to the deformation-induced increased Ca^2+^ permeability in on cell patch clamp experiments [19, 20], but it was not clear whether this was an incidental association of relevance to lifespan dehydration. With PzA set to zero in the lifespan model, dehydration was severely reduced (Fig 4, curve 5) showing an extremely limiting effect of the anion permeability, much more powerful than the one predicted for single transits [3]. These results show that increased anion permeability is a necessary companion to PzCa in the generation of the lifespan pattern (Fig 4, curve 1). The attribution of the increased PzA to PIEZO1 in the simulations is purely operational; its actual mediation remains an open question [20]. The model simulations revealed an additional unexpected role of the increased anion conductance, as shown by comparing curve 7 with curves 5 and 6 in Fig 4. If PzA was not increased, whether or not in association with increased PzCa, the relatively rapid initial dehydration phase was absent. This result and additional tests showed that the relatively rapid early dehydration phase is largely PzA-dependent.

### Effects of PIEZO1-mediated sodium and potassium permeabilities

In sharp contrast to the powerful effects of PzCa and PzA on the kinetic pattern of the dehydration response, increasing PzNa and PzK from zero to over two orders of magnitude above ground sodium and potassium permeabilities had minor effects (Fig 4, curves 2 and 3), mostly on the timing and magnitude of the late density reversal response. The minor effects of large increases in PzNa and PzK render the Lifespan model a poor guide on the likely values of real PIEZO1-mediated Na^+^ and K^+^ permeabilities during capillary transits, the reason for setting their default values to zero in our minimalistic approach to the design of the reference curve.

### Analysis of the effects of pump decay rates and Gardos channel inhibition

Blocking the Gardos channel suppresses fully the dehydration-densification component of the response (Fig 5, curve 6) stressing the critical, albeit indirect role of this channel in the PIEZO1-mediated lifespan densification process. Gardos channel activity has been shown to marginally decline with RBC age [13]. Model tests of declining Gardos Fmax within observed boundaries had no detectable effects (not shown).

By setting the Na/K pump decay to start from the beginning (TNaP = 0) the dehydration rate becomes substantially reduced whether late reversal is curtailed (Fig 5, curve 3) or preserved (Fig 5, curve 4). Multiple additional simulations testing different pump-decay rates showed that early Na/K pump inhibition, by allowing progressive net NaCl gains partially offsets the dehydrating-densifying effects of Gardos channel mediated KCl losses, keeping the dehydration profile below observed boundaries. This explains the requirement of a substantial delay in the onset of Na/K pump decay for the reference patterns to emerge (Fig 5, curve 1), and suggests an evolutionary component in the development of a delayed Na/K pump decay and in the prevention of significant early decay. These simulations, of course, do not rule out some minor level of early pump decay; they only stress the absolute requirement of late decay for density reversal.

The rate of decline in Na/K pump activity found to approximate the observed extents of density reversal predict a decline in Na/K pump sodium efflux from about −3.2 mmol/Loch on day 80 to −2.5 mmol/Loch on day 120, a 22% flux decline (data from reference csv file). The real decline in pump Fmax, on the other hand, set at e^-kt^, with k=3•10^−5^ (min^-1^), is 82% within this 40 day period. The reason for the discrepancy is that the flux was being strongly stimulated by the increasing intracellular sodium concentration (Fig 6), in turn the result of sharp progressive Fmax pump decline. Therefore, flux comparisons performed without correcting for differences in intracellular sodium concentrations between RBCs of different ages would tend to underestimate the true magnitude of the Na pump decay, as noted by Cohen et al., [15].

Curves 2 and 5 in Fig 5 illustrate the powerful and extremely sensitive effects of a faster (Fig 5, curve 2) or slower (Fig 5, curve 5) PMCA decay rate relative to the reference curve 1, and illustrate the dominant role of PMCA decay in the control of the densification stage throughout most of the lifespan period, the inflexion reflecting the accelerating effects of an exponentially-defined decay process. The conditions represented by curves 2, 3 and 8 in Fig 4 and by curve 2 in Fig 5 are equal to curves 1 in their entitlements to represent compliant densification patterns.

The interesting insight arising from this analysis is that declines in pump activity, generally assumed to result from cumulative protein damage in the absence of biosynthetic renewal, may have an evolutionary biased component in their magnitude and timing.

### Predicted lifespan changes in intracellular Na^+^, K^+^ and A^-^ concentrations

Over the first 70-80 days, the model predicts a slow quasi linear fall in CK in excess of the much slower CNa rise, with the CA decline following the trend set by the sum of the net cation concentration changes (Fig 6). After about day 80 the trend rapidly reverses causing a delayed CA reversal as soon as sodium gains exceed potassium loses. These model-predicted trends are in full agreement with all previous measurements of age-attributed sodium and potassium content changes in RBCs.

### Predicted changes in intracellular pH (pHi) and proton ion concentration (CH) and their relation to the parallel changes in intracellular concentrations of diffusible anions (CA)

The kinetics of the pHi changes follows closely that of the change in intracellular anion concentration (Fig 7A) as expected from an anion ratio driven process. As explained in detail before [2], the Jacob-Stewart mechanism (JS) drives the rH ratio ([H^+^]_i_/[H^+^]_o_) to match the rA ratio ([A^-^]_o_/[A^-^]_i_), a match reflected in the mirror images of the CH and CA curves of Fig 7B. The large capacity of the JS mechanism ensures that the infinitesimal changes in anion concentration ratio generated during each transit are rapidly approximated by the proton concentration ratio during inter-transit periods. Here again, isolated measurements of pHi in density-segregated RBCs fully support gradual cell acidification as predicted by the model (Fig 7A).

### Statistical compliance of predicted volume changes with cell age

The default settings of the csv-file outputs in the lifespan model were designed to report changes in variable values at daily intervals. Each row in the csv file reports the values predicted for each variable for a particular day. The condition of the RBC at time = 0 is defined by the entries in the Reference State. Each column reports how each variable changes daily throughout the 120 days of the modelled lifespan offering a distribution of values open to statistical comparisons with measurements. Measured variations in RBC properties combine production line diversity, as RBCs emerge from the bone marrow, with changes caused by circulatory aging. Model-predicted variations, on the other hand, report only variations caused by circulatory aging of a single cell defined with the default “birth” properties at the Reference State. It follows that the predicted coefficient of variation of the columns representing a single day cohort, by lacking the production line variation component, has to be smaller than that of the corresponding measured variable in real samples. The haematological variables for which we have extensive and reliable measurements are RBC volume, haemoglobin contents and haemoglobin concentration [8]. Of these, the only one allowing comparison between predicted and measured values is RBC volume, because the haemoglobin content of the model RBC, fixed in the Reference State, is assumed to remain constant throughout the RBC lifespan. The mean and standard deviation values of the relative cell volume distribution for the cell represented by the reference pattern (Fig 3) were 0.956 and 0.069, respectively, rendering a coefficient of variation of 7.2%, well below measured values around 12-13%. To the extent these predicted values approximate reality they suggest that about half the volume variations in measured RBC samples are the result of aging-induced changes in RBC hydration state.

### Hyperdense collapse and PIEZO1-mediated calcium permeability

A relatively minor reduction in PzCa, from 70/h to 60/h (Fig 4, curve 4) proved sufficient to prevent dehydration reaching the documented range. On the other hand, increasing PzCa above 70/h eventually overwhelmed the declining Ca^2+^ extrusion capacity of the PMCA and triggered a sharp hyper-dense collapse, an extreme dehydration response (Fig 8A). The model predicts partial volume and density recovery post-collapse (Fig 8A) resulting from net NaCl and fluid gains following progressive Na/K pump decay, suggesting a potential for full lifespan survival of the cells (Figs 8A-B). It remains an intriguing open question whether such a sequence may, very occasionally, occur in vivo. Volume collapse takes a few days to evolve and much longer to recover so that the probability to bypass spleen sinusoids during the hyperdense stages seems remote.

The results in Figs 4 and 8 show that OS*PzCa values navigate narrow ranges of variation between failing to densify sufficiently and hyperdense collapse. The 70h^-1^ PzCa value at an OS of 0.4s used for the Ref simulations (Fig 3) represents an increase of around three orders of magnitude over the basal calcium permeability of the RBC membrane, of 0.05h^-1^, the value found to fit the observed physiological rates of PMCA pump-leak turnover [21]. In more comparable permeability units this represents an effective increase in RBC membrane calcium permeability from ∼10^−9^cm/s, one of the lowest in nature, to 10^−6^cm/s. The model-predicted narrow range of OS*PzCa values represents a complex mix combining OS duration, number of PIEZO1 channels per cell and the intrinsic Ca^2+^ permeability of each channel as influenced by cell and medium sodium and potassium concentrations [22]. In vivo, some of these components will vary from transit to transit, OS with speed of flow, and number of PIEZO1 channels activated by extent of deformation. Ultimately, according to the model, it is the aggregate of Ca^2+^ loads that PIEZO1 activation induces during each capillary transit ([3]; Fig 5), averaged over many transits, what counts for shaping the Lifespan pattern.

### Effects of RBC volume fraction during capillary transits on Lifespan patterns

The biphasic volume change patterns of RBCs during single capillary transits proved to be almost identical when simulated with cell volume fractions (CVFs) of 0.9 during their brief capillary transits, or of 0.00001 throughout both transit and inter-transit periods [3]. However, when the effects of the two alternative transit CVF values were compared on the lifespan model, under identical conditions and parameter values, the red cell volume of the low-CVF cell slowly drifted towards a late hyperdense collapse (Fig 9, red). At high CVFs the transfer of calcium from medium to cells was bound to decrease the medium calcium concentration significantly more than at low CVFs. Cell calcium gains during capillary transits at low CVFs were therefore expected to be slightly larger than those at high CVFs under comparable condition. These differences were too insignificant to cause noticeable volume changes on single transits [3], but had the potential to accumulate and overwhelm the Ca^2+^ extrusion capacity of a PMCA decaying at the REF-set rate when balancing the calcium gains of a low-CVF cell, as proved to be the case (Fig 9). The inset of Fig 9 shows the expected trend of change in [Ca^2+^]_i_ as recorded at the end of the last capillary transit each day for low-CVF conditions (red) and for REF conditions (black). The restoration of comparability between high- and low-CVF conditions when the rate of PMCA decay at low-CVF is reduced (Fig 9, green) confirms that the discrepant volume response can be entirely attributed to differences in cumulative calcium gains. This outcome then justifies the choices made of modelling capillary transits and lifespans with the more realistic high-low CVF transitions between capillary and systemic circulation using the Restore Medium subroutine [3].

### Iirreversibly sickled cells (ISCs): fast track lifespan, hyperdense collapse, and terminal density reversal

The condition of the collapsed cells resembles that of the subpopulation of sickle cells known as irreversibly sickled cells (ISCs) [23-25]. These cells originate from a subpopulation of stress reticulocytes, have an extremely short, 4-7 day lifespan, dehydrate rapidly by a calcium-dependent fast-track mechanism on egress from the bone marrow [26, 27], persist in the systemic circulation for a few days in a hyperdense condition, terminally gaining NaCl and rehydrating back to a low density state [9]. In their hyperdense state, ISCs are responsible for vaso-occlusion leading to multiple organ infarctions, pain crisis, and for most of the clinical symptoms of sickle cell disease (SCD) irrespective of their proportion in the circulation [28, 29]. Early spleen infarction in infancy leads to functional asplenia [30] and prevents effective clearance of ISCs in SCD patients.

The potential for hyperdense collapse exposed by the lifespan model prompted a search for conditions which could emulate the circulatory trajectory of ISCs. Figure 10 shows a predicted lifespan for a sample seven-day ISC. The open state attributed to PIEZO1 as Psickle [31], the permeability pathway generated by the interaction of deoxy-haemoglobin S polymers with the inner membrane surface, was set at 30 seconds emulating the observed extended periods of elevated calcium permeability in deoxygenated conditions that might be expected during inter-transit periods in the venous systemic circulation [32]. Sickling follows a ∼ 40^th^ power dependence on the concentration of deoxy-haemoglobin S (deoxy-HbS) [33, 34]. Yet sickle reticulocytes, still far short of the full HbS complement of the matured red cell, and with a far lower haemoglobin concentration, showed a sharp and reversible permeabilization response to K^+^, Na^+^ and Ca^2+^ on deoxygenation [26].

**Figure 10.**
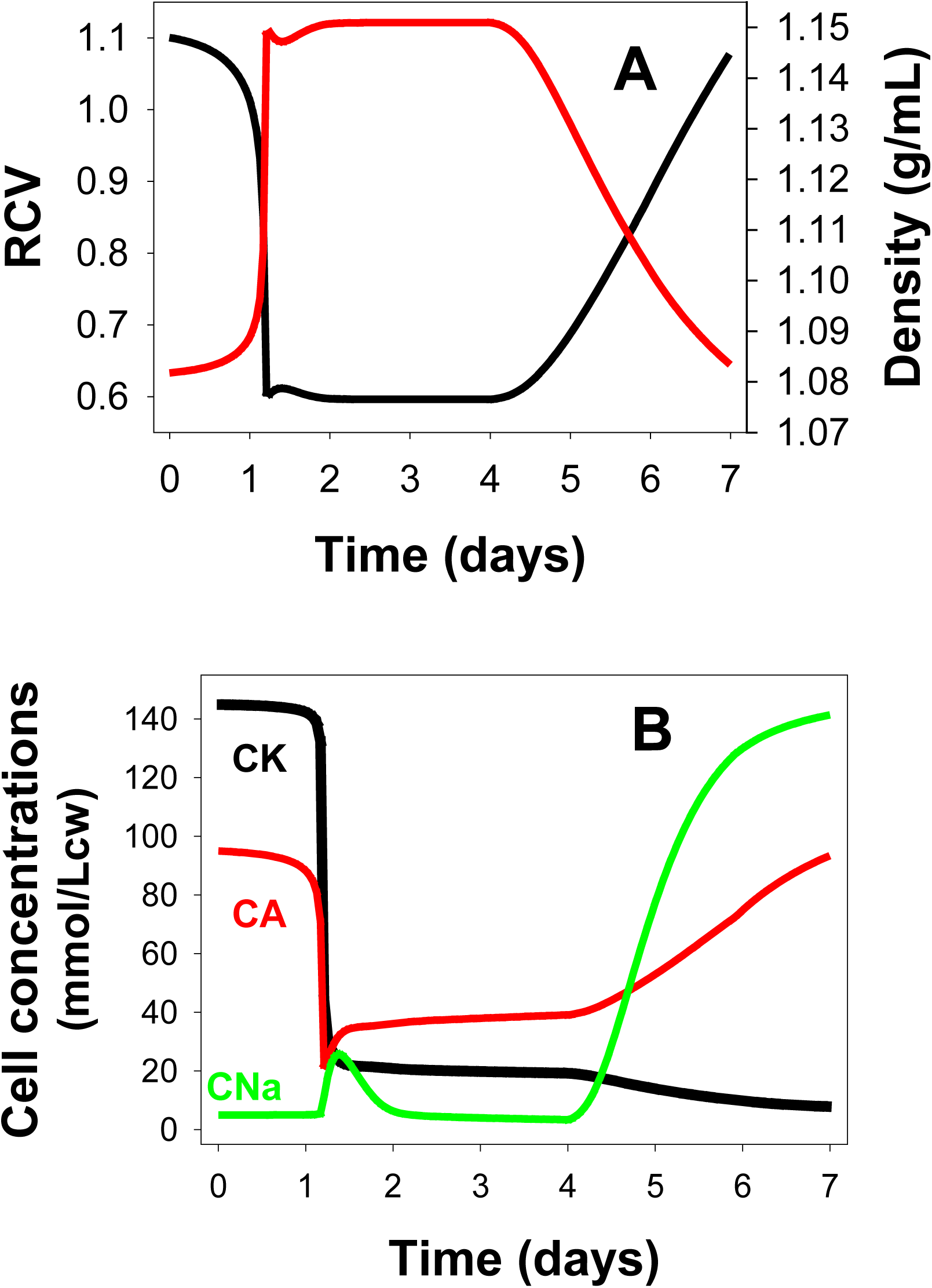
Simulating the lifespan of a seven day irreversible sickle cell (ISC), showing the predicted changes in RCV (black) and density (red), (panel A), and in the cell concentrations (panel B) of K(CK), of Na(CNa), and of diffusible anion(CA). The parameters used were: Lifespan duration, 168h (7 days); Data output periodicity: 60m; kNaP, 2e-3/m; kCaP, 5e-5/m; TNaP, 5760m (4 days); PIEZO1 open state, 30s; PzCa, 5.7/h; PzA, 50/h; PzNa, 0; PzK, 0.

With a 30s open state, a marginal PzCa increase to 5.7/h was sufficient to induce hyperdense collapse within just over a day, as expected for a fast-track dehydration mechanism (Fig 10A) [25]. Delayed Na/K pump decay initiated rapid rehydration of the ISCs after three days in hyperdense state, rehydration induced by late NaCl and fluid gains (Fig 10B), as observed [9]. The rapid Na-K gradient and density reversal of ISCs opens the question of whether intravascular osmotic lysis or erythrophagocytosis by immune exposure seals the terminal lifespan of this subpopulation of sickle cells. Allowing for its limitations, the current simulation shows the power of the same membrane components and capillary transit dynamics used to predict the lifetime homeostatic changes of normal, HbA RBCs to emulate the essentials of ISC generation. It is hoped this preliminary result will stimulate further investigations with an updated model representation of sickle cell reticulocytes [26], including the documented contribution of the KCC3 K:Cl cotransporter during the rapid dehydration-acidification stage [26, 35-37].

### Spontaneous inactivation of PIEZO1 or Psickle channels

The 30s open state attributed to Psickle in the simulation of Fig 10 was meant to represent the active condition of this channel during inter-transits in the venous systemic circulation, implying a condition of suspended or delayed inactivation supported by a variety of experimental observations. RBCs from heterozygote HbSA subjects with largely normal membrane properties, when exposed to deoxy-sickling pulses lasting hours remained in a high Ca^2+^ permeability state for the full duration of the pulses [38]. This was also shown to be the response of discocyte-stage HbS RBCs during extended sickling periods in vitro [32]. Thus, inactivation appears to be suspended for the duration of the sickled condition. When PIEZO1 channel activation is elicited in HbA RBCs by contact with electrodes, inactivation also appears disrupted or profoundly delayed [19, 39]. On the other hand, calcium signals recorded from Fluo-4-loaded RBCs traversing microfluidic constrictions under pressure regimes that ensure transit times approximating those in vivo, conditions assumed to preserve normal membrane-cytoskeletal connectivity, showed reversible fluo-4 signals reporting elevated [Ca^2+^]_i_ only during transits, the expected response of PIEZO1 channels with preserved spontaneous inactivation ([39] their figure 3).

Taken together, these disparate observations suggest that local disruption of the normal structural connections between the plasma membrane and cytoskeleton, either by HbS polymers protruding through the cytoskeletal mesh or by electrode contact or suction, affect some or all of the Psickle or PIEZO1 channels expressed in the RBC membrane [40] somehow disturbing configurations required for spontaneous inactivation functionality. Such a mechanism may also hold the answer to the unexplained nature of the high cation leak pathways of reticulocytes.

The mean Ca^2+^ and Na/K pump-leak turnover of reticulocytes is about ten-fold higher than that of mature RBCs [41, 42]. The nature of the Ca^2+^, K^+^ and Na^+^ leak pathways remains a mystery [26, 27, 37]. An intriguing possibility worth future investigation is that amidst the profound cytoskeletal-membrane remodelling processes taking place after erythroid cell enucleation the high cation leaks are mediated by open PIEZO1 channels, roughly estimated in the hundreds per cell at the reticulocyte stage [40]. As spontaneous inactivation functionality becomes established with progress towards the membrane-cytoskeletal configuration of mature RBCs so does the declining magnitude of the leak fluxes.

These considerations highlight a set of open questions arising from the current lifespan investigation implicating PIEZO1 inactivation as the target of interest in pathologies affecting RBC hydration and suggest altered membrane-cytoskeletal interactions as the focus of attention for explaining the molecular mechanism of disrupted PIEZO1 inactivation.

## Discussion

We approached the investigation of the mechanisms shaping the lifespan changes of human RBCs, a subject inaccessible to direct experimentation, by applying the mathematical-computational model of human RBC homeostasis introduced in the first paper of this series [2]. We started by questioning the nature and range of RBC responses to be expected from deformation-induced PIEZO1 activation during single capillary transits [3], and followed this up in this paper with a systematic exploration of the dynamic combinations of homeostatic processes that could deliver the documented patterns of change throughout the ∼2•10^5^ transits RBCs experience during their long lifespan using a model extension adapted for lifespan studies (Fig 1). Initial apprehensions when facing the large undocumented parameter space during preliminary tests were soon allied by the powerful constraining effects of fitting the well established lifespan sequence of prolonged densification and late reversal, exercise interested readers are encouraged to emulate and explore further.

A first result from the lifespan model simulations was the demonstration of the inadequacy of repeated capillary transits to generate long-term progressive RBC densification on their own (Fig 2), their cumulative power fading rapidly within days to minimal densification levels, thus ruling out the quantal hypothesis [43, 44]. The mechanism that finally emerged involved a complex interplay among a quartet of membrane transporters (PIEZO1 and Gardos channels, PMCA and Na/K pumps) involving a tightly defined decay patterns for the pumps (Figs 3-5) and additional modulating influences by all other membrane and homeostatic components of the RBC (Figs 6-8), reported and analysed in detail in Results and Analysis.

Looking back at the variety of conditions which failed on compliance with the established densification-late-reversal pattern (Figs 4-5), there are clear indications that RBC volume stability and hence adequate rheological performance throughout extended lifespans could also be attained by other, apparently simpler alternatives. Playing with the model unconstrained by facts, one alternative emerged which, surprisingly, was also tightly constrained, that of a RBC without PIEZO1 channels (PzX = 0) and without Gardos channels (PKGardosMax = 0 in RS), but with a well balanced pattern of pump decays (in min^-1^, kCaP = 8*10^−5^; kNaP = 1*10^−6)^), a simple pump-controlled duet mechanism in which the opposite swelling-shrinking effects elicited by decaying Na/K and PMCA pumps, respectively, remain well balanced throughout. A RBC like this, free from sudden permeability changes during capillary transits by the absence of PIEZO1 channels, and exempt from hyperdense collapse threats (Fig 8) by the absence of Gardos channels could sustain excellent volume stability and optimal rheology throughout extended lifespans at slightly lower metabolic cost that a quartet cell exposed to periodic PMCA stimulation. This prompted the question of what favoured or determined the evolution of the quartet mechanism in human RBCs.

There is no strong argument for a selective preference of quartet over duet mechanisms or other alternatives. In different species the universals of optimal economy and rheology providing extended lifespans are realized with very different strategies, typical of adaptive solutions on the go operating on pre-existing conditions [45]. There are well documented instances of species whose RBCs lack Gardos channels [46], have kinetically diverse and even absent Na/K pumps [47, 48] and varied Na/K concentration ratios [49], have different constellations of membrane transporters controlling RBC volume and homeostasis [50], differ substantially between foetal and mature RBCs [51], have completely different cytoskeletal structures, with and without vestigial organelle retention [52, 53]. There is no information available yet on how widespread the presence of PIEZO1 channels is in RBCs from different species, and therefore on how central its role may be in the dynamics of capillary circulation. So far, the only documented constant in all species appears to be the calcium pump around which all the different lifespan strategies of RBCs evolved.

Mutant PIEZO1 channels in hereditary xerocytosis (HX) were found to exhibit a number of kinetic abnormalities the most prominent of which was a marginally reduced inactivation kinetics following brief stretch-activation pulses [54]. Simulations with the Lifespan model show how relatively small inactivation delays can lead, after myriad capillary transits, to profound RBC dehydration approaching hyperdense collapse (Fig 8) with similarities to observed alterations in HX RBCs [55]. Protection against severe falciparum malaria, the reason for the persistence of many genetic mutations affecting RBC hydration in human populations, is contributed by two common conditions: anaemia and the presence of subpopulations of dense RBCs which falciparum merozoites fail to infect [56], thus preventing the build up of the high parasitaemias required to cause cerebral malaria, the main malaria killer [57, 58].

The lifespan model opens the way for further in depth studies on the changes in RBC homeostasis during circulatory senescence by further exploring the effects of alternating oxy-deoxy capillary transits. With growing information databases on the genetics and pathologies associated with the transport systems that control the lifespan of RBCs, model versions encoding known or hypothesized abnormalities of those transporters may become useful tools in furthering the understanding of the pathophysiology and clinical manifestations of the diseased conditions. Within the mathematical-computational framework of the red cell and lifespan models applied here for human RBCs, components can easily be modified and adapted to explore RBC homeostasis, circulatory dynamics and lifespan strategies across species paving the way for future studies on the comparative circulatory biology and pathology of RBCs.

## Acknowledgements

The background research work on which this investigation was based was supported over four decades by funding agencies in the UK and in the USA (VLL PI or co-PI). **In the UK:** Biotechnology and Biological Sciences Research Council (BB/E008542/1; BB/F001630/1, BB/F001673/1, and BB/H024867/1), Engineering and Physical Sciences Research Council (EP/E059384); The Wellcome Trust (064124; 061269; 059725; 030699; 033876; 15543; 17358; 13056), and The Medical Research Council (G8211073CA). **In the US:** NIH 2-RO1 HL28018-19; 2-RO1 HL21016-11. The authors are grateful to Serge L. Y. Thomas and Daniel J. Lew for helpful comments and suggestions on the material contained in the three manuscripts of this series.

